# Structural covariance of amygdala subregions is associated with trait aggression and endogenous testosterone in healthy individuals

**DOI:** 10.1101/2021.07.09.451771

**Authors:** Martin Göttlich, Macià Buades-Rotger, Juliana Wiechert, Frederike Beyer, Ulrike M. Krämer

## Abstract

Many studies point toward volume reductions in the amygdala as a potential neurostructural marker for trait aggression. However, most of these findings stem from clinical samples, rendering unclear whether the findings generalize to non-clinical populations. Furthermore, the notion of neural networks suggests that interregional correlations in grey matter volume (i.e., structural covariance) can explain individual differences in aggressive behavior beyond local univariate associations. Here, we tested whether structural covariance between amygdala subregions and the rest of the brain is associated with self-reported aggression in a large sample of healthy young students (n=263; 51% women). Salivary testosterone concentrations were measured for a subset of n=76 participants (45% women), allowing us to investigate the influence of endogenous testosterone on structural covariance. Aggressive individuals showed enhanced covariance between superficial amygdala (SFA) and dorsal anterior insula (dAI), but lower covariance between laterobasal amygdala (LBA) and dorsolateral prefrontal cortex (dlPFC). These structural patterns overlap with functional networks involved in the genesis and regulation of aggressive behavior, respectively. With increasing endogenous testosterone, we observed stronger structural covariance between centromedial amygdala (CMA) and medial prefrontal cortex in men and between CMA and orbitofrontal cortex in women. These results speak for structural covariance of amygdala subregions as a robust correlate of trait aggression in healthy individuals. Moreover, regions that showed structural covariance with the amygdala modulated by either testosterone or aggression did not overlap, speaking for a more complex role of testosterone in human social behavior rather than the simple assumption that testosterone only increases aggressiveness.

## Introduction

Aggressive behavior, often defined as “any form of behavior directed towards the goal of harming or injuring another living being who is motivated to avoid such treatment” (Baron and Richardson, 1994), remains of utmost societal concern (Hoeffler, 2017). Finding reliable predictors of aggression is thus an important goal that is yet to be fulfilled. Because metrics of brain structure remain remarkably stable over time in adults (Elliott et al., 2019), they hold great promise as a marker of interindividual differences in aggressiveness. Aggressive behavior is thought to depend on a subcortical circuit comprising the amygdala, ventromedial hypothalamus, and periaqueductal gray, which are all subject to prefrontal control (Nelson and Trainor, 2007). Among these structures, lower amygdala volume has been consistently associated with increased self-reported aggression across a range of clinical populations (Driessen et al., 2000; Matthies et al., 2012; Pardini et al., 2014; Rogers and De Brito, 2016; Thijssen et al., 2015; Yang et al., 2009; Zhang et al., 2013). The link between aggressiveness and low amygdala volume is perhaps the most replicable finding in this subfield, with few exceptions (e.g. Mancke et al., 2018; Varkevisser et al., 2020).

Gray matter reductions linked with aggressive behavior are also apparent in other regions of the aggression circuitry. An early study on this topic found that individuals with pathological aggression have reductions of around 11% in prefrontal cortex volume relative to healthy controls and substance-dependent subjects (Raine et al., 2000). Around the same time it was reported that aggressive epilepsy patients also evince reductions in prefrontal volume as compared to controls and non-aggressive patients (Woermann et al., 2000). More recent investigations have narrowed down the focus to specific regions of the PFC, suggesting that volume reductions in pathologically aggressive individuals are most pronounced in orbitofrontal and dorsolateral prefrontal portions (Leutgeb et al., 2016; Raine et al., 2011; Tiihonen et al., 2008). Furthermore, volume in these areas has been negatively associated with both self- and clinician-rated antisocial behavior among patients with antisocial personality disorder (Raine et al., 2011). Other research has shown that the volume of the striatum, a brain region involved in reward processing and action initiation (Balleine et al., 2007), is associated with greater aggression in children (Ducharme et al., 2011), violent offenders (Schiffer et al., 2011) and psychopaths (Glenn et al., 2010).

This brief overview of the literature indicates that aggressiveness is associated with reduced gray matter in amygdala, orbitofrontal cortex, and dorsolateral prefrontal cortex but relatively larger striatal volume. It is also apparent, however, that most of these studies have been performed in clinical or forensic samples. Despite the inherent value of such investigations, some characteristics of these samples (e.g., comorbidity, long-term treatment) make it difficult to draw clear conclusions from the observed brain-behavior associations. Furthermore, sample sizes in studies from hard-to-reach populations are often low, which is likely to yield imprecise estimates of effect size (Maxwell, 2004). These limitations call for studies with larger samples of healthy individuals, as they might help to ascertain which neurostructural correlates of aggression are dimensional in nature and which ones are disorder specific. For instance, the lower ventral prefrontal volume often observed in aggressive clinical samples has been recently reported in healthy subjects with high trait physical aggression (Chester et al., 2017).

Another feature of many volumetric studies on the topic is the focus on mass univariate associations between gray matter structure and aggressive behavior. Although informative, this traditional approach fails to capture covariation between brain areas (i.e. structural covariance) in the same manner that local estimates of hemodynamic activity do not directly measure connectivity between distant regions (Camara et al., 2009). Structural covariance is however a well-documented phenomenon thought to reflect joint development and/or sustained co-activation. Indeed, correlated structural patterns arise during childhood, overlap with intrinsic functional networks, and break down in the presence of neurodegeneration (Alexander-Bloch et al., 2013; Seeley et al., 2009; Zielinski et al., 2010). Of note, a developmental study suggested that the relationship between testosterone and aggression is partly dependent on covariance between amygdala and medial prefrontal cortex (Nguyen et al., 2016). Investigating covariation between distal brain areas might thus reveal neural network markers of aggression that standard methods might have missed out on.

A particularly relevant question in this regard is whether specific amygdala nuclei show divergent patterns of covariance with other brain areas, and whether this is linked with aggressive behavior. Although the amygdala as a whole is consistently associated with aggressive behavior in humans (Fanning et al., 2017), some of its subregions carry out partly opposing roles in orchestrating aggression. Specifically, both the centromedial (CMA) (Han et al., 2017; Nordman et al., 2020) and the superficial amygdala (SFA) (Yamaguchi et al., 2020; Zha et al., 2020) are thought to promote aggressive responses to threat via their numerous projections to the periaqueductal gray and other brainstem nuclei (Abivardi and Bach, 2017; Han et al., 2017; Saygin et al., 2011). This is achieved by relying on ventrolateral aspects of the ventromedial hypothalamus and the bed nucleus of the stria terminalis (Nelson and Trainor, 2007; Nordman et al., 2020). Interestingly, there is some evidence that aggression-related alterations in amygdala volume are circumscribed to subregions broadly corresponding to the central and superficial nuclei (Gopal et al., 2013). The laterobasal amygdala (LBA), in contrast, inhibits centromedial input to the brainstem and thereby facilitates active, flexible coping (Terburg et al., 2018). In line with this assumption, LBA activity has been linked with both avoidance (Buades-Rotger et al., 2017) and retaliation (Buades-Rotger et al., 2016b) in response to interpersonal provocation. Among all amygdala nuclei, the LBA receives the highest density of axonal terminals originating from the orbitofrontal cortex (Ghashghaei and Barbas, 2002), which is thought to provide an additional top-down, aggression-regulatory input to the CMA and SFA (Nelson and Trainor, 2007) through its direct connections with these structures (Abivardi and Bach, 2017). Further, compared with the SFA/CMA, the LBA has stronger anatomical connectivity with the anterior temporal pole (Bach et al., 2011), an empathy-related brain area which shows volume reductions in individuals with high levels of reactive aggression (Bertsch et al., 2013). The existing evidence hence suggests that dorsal-posterior aspects of the amygdala promote aggressive behavior via downstream connections with other subcortical structures, whereas the LBA provides modulatory input.

In sum, most volumetric studies on aggression have employed small, clinical samples and have not tested for covariance patterns between different brain areas. Moreover, the structural networks of the main amygdala nuclei groups remain to be delineated, which might help to clarify discrepancies regarding the role of each region in aggression. To address these issues, we tested whether structural covariance between amygdala subregions and the rest of the brain is associated with self-reported aggressive behavior in a large sample (n=263) of healthy young students.

Finally, as Nguyen et al. (2016) found that covariation between amygdala and medial prefrontal cortex mediated the positive relationship between testosterone and aggression in a sample composed mostly of children and adolescents, we set out to extend this research to young adults in whom brain maturation is expected to be completed. Salivary testosterone concentrations were available for a subset of n=76 students (45% women), allowing us to investigate the influence of endogenous testosterone on the covariance between amygdala and voxel-wise gray matter volume. Medial prefrontal cortex and the orbitofrontal cortex belong to the brain regions with high concentrations of androgen receptors (Cunningham et al., 2012; Finley and Kritzer, 1999; Tobiansky et al., 2018). We thus aimed to test whether aggression- and testosterone-related structural covariance converge on the same amygdala-dependent networks, providing further support for the hypothesis of Nguyen et al. (2016).

## Methods

### Participants

The final sample comprised 263 healthy young students (age=23.28±2.80 years, 129 women). All participants were free of psychiatric and neurological disorders (self-report) and all T1-weighted images were inspected by an experienced neuroradiologist. The study was designed in accordance with the Declaration of Helsinki and approved by the Ethics Committee of the University of Lübeck prior to recruitment. Written informed consent was obtained from all participants. The structural data had been collected as part of different studies, whose functional data have partly been reported previously (Beyer et al., 2017; Beyer et al., 2013; Beyer et al., 2015; Beyer et al., 2014a; Beyer et al., 2014b; Buades-Rotger et al., 2017; Buades-Rotger et al., 2016a; Buades-Rotger et al., 2016b; Buades-Rotger et al., 2016c).

### Trait Aggression Measure

As a measure of trait aggression, all participants filled out a German version of the Buss and Perry Aggression Questionnaire (AQ) (Buss and Perry, 1992; Herzberg, 2003), which comprises four subscales: anger, physical aggression, verbal aggression and hostility. Answers in the AQ are given on a 5-point Likert scale, whereby higher values mean higher aggressiveness.

### Testosterone

Participants provided saliva samples in plastic vials (women: 4 ml Cryovials from Salimetrics, men: SafeSeal micro tube 2ml from Sarstedt) with the passive drooling technique before and after scanning. We froze these samples at −20°C and shipped them in dry ice to a reference laboratory in Manchester (UK) for analysis once the study was completed. Free (i.e., unbound) testosterone concentrations were estimated with liquid chromatography tandem mass spectrometry (LC-MS/MS) as described elsewhere (Keevil et al., 2014). Intra- and inter-assay coefficients of variation have been reported to be 5.3% and 9%, respectively, with a lower limit of quantification of 5 pmol/L (Keevil et al., 2014).

### Data acquisition

We recorded high resolution T1-weighted images from all subjects. Three data sets, which were acquired at three different scanners, were included in the analysis. The first data set (n=142 subjects) was obtained at a 3T Philips Achieva MRI scanner (Philips Healthcare, the Netherlands) equipped with an eight-channel head coil applying a T1-weighted 3D turbo gradient echo sequence with SENSE (matrix 170×240×240; spatial resolution 1×1×1 mm^3^, repetition time 8.2 ms; echo time 3.8 ms; flip angle 8°). The second data set (n=74) was recorded at a Philips Ingenia 3T MRI scanner applying a T1-weighted 3D turbo gradient echo sequence (matrix 170×240×240; spatial resolution 1×1×1 mm^3^, repetition time 7.8 ms; echo time 3.5 ms; flip angle 8°). The third data set (n=47) was recorded at the CBBM Core Facility Magnetic Resonance Imaging using a 3-T Siemens Magnetom Skyra scanner equipped with a 64-channel head-coil. Structural images of the whole brain were acquired using a 3D T1-weighted MP-RAGE sequence (repetition time 2300 ms; echo time 3.0 ms; inversion time 900 ms; flip angle 9°; 1×1×1 mm^3^ resolution; matrix 192×320×320).

### Analysis

We performed standard image preprocessing for voxel-based morphometry (VBM) using the CAT 12 toolbox (Computational Anatomy Toolbox 12 for SPM; http://dbm.neuro.uni-jena.de/cat/). The T1-weighted images were normalized to the MNI template space and segmented into gray matter, white matter and cerebrospinal fluid applying the updated segmentation algorithm implemented in SPM12 (University College London, Wellcome Trust Centre for Neuroimaging, http://www.fil.ion.ucl.ac.uk/spm/). Spatial normalization was performed to achieve voxel-wise correspondence across subjects. The produced tissue class images were resampled to a voxel size of 1.5×1.5×1.5 mm^3^, bias corrected for intensity non-uniformities and modulated, i.e., the voxel values were scaled by the amount of volume changes derived from the spatial normalization. This allows comparing the absolute amount of gray matter (‘gray matter volume’) across subjects. The modulated gray matter images were spatially smoothed by a Gaussian kernel with a FWHM of 8mm.

We performed data quality checks including image noise, bias, and the voxel-wise mean correlation between gray matter volumes across subjects. For each pair of subjects, the correlation between their gray matter volumes was calculated. The mean of the Fisher z-transformed correlation coefficients for each individual subject was then used as a data quality indicator. We visually inspected all images where one of the quality indictors differed from the mean by more than two standard deviations. Fifteen subjects showed considerable motion artefacts and were excluded from the analysis resulting in n=263 subjects in the final data sample.

We set up a multiple regression analysis using SPM12. The independent variables were amygdala subregion volume, self-reported trait aggression (AQ score), and crucially their interaction. We also included age, gender, and scanner model as covariates in the analysis. A global scaling to the intracranial volume (TIV) was applied. Only voxels with a gray matter probability larger than p=0.1 were included in the analysis. We defined the amygdala seeds using the Anatomy toolbox (Eickhoff et al., 2007) which provides a parcellation of the amygdala into three distinct subregions that differ in their structure, function and connectivity profile (Bzdok et al., 2013): laterobasal (LBA), centromedial (CM) and superficial (SF) nuclei group. T-maps were calculated for the two main effects and their interaction. The statistical images were assessed for cluster-wise significance using a cluster-defining threshold of p=0.001. A topological family wise error (FWE) procedure was used to correct for multiple comparisons. The p<0.05 FWE corrected critical cluster size was k=734 voxels. We applied the same approach to investigate the effect of testosterone on brain structure and amygdala covariance. For analyses involving testosterone, we decided against combining the male and female data sets as they were recorded with different MRI scanners which means that effects of gender and confounding scanner effects cannot be disentangled.

A positive or a negative interaction term only carries the information about whether the correlation between amygdala and target region volume is increasing or decreasing with increasing aggression. However, for the interpretation of the results it is important to know whether the correlation is positive or negative for different subgroups with similar aggression scores. We therefore examined the interactions in more detail. For each subject, we extracted the mean gray matter volume in the amygdala subregions (LBA, CMA and SFA) and in the clusters showing a significant interaction between amygdala volume and AQ score and performed a regression to remove variance explained by the total intracranial volume, gender, age, and scanner model in each region. We then subdivided our data sample into three equally large subsets of increasing aggression score (or testosterone level) and calculated the correlation between the volume in the amygdala seed region and the target region. It should be stressed that this is a qualitative analysis which guides the interpretation of the results and which allows us to explore whether they hold for different subscales of the AQ or for subsamples of the whole data set, i.e., different scanner type or gender. Statistical tests for differences between AQ or testosterone subgroups are not valid due to a selection effect.

## Results

### Demographic data and testosterone level results

We summarized all relevant demographic, behavioral and physiological information in Table 1. The mean testosterone concentration in the third dataset (Skyra) was 212.3±63.5 pmol/l (n=44 subjects). Testosterone levels for 37 female subjects in the second dataset (Ingenia) were available. The mean testosterone level in that group was 16.9±6.9 pmol/l. The testosterone concentrations are in good agreement with expected values for male and female subjects in that age range (Keevil et al., 2014). We found a positive correlation between testosterone levels and physical aggression in male subjects: Pearson’s □=0.37 and p=0.020. We applied a z-transformation to the testosterone concentrations for both groups of male and female subjects to combine the two data samples. In the combined data set we found a significant positive correlation between aggression (AQ score) and testosterone: □=0.23 and p=0.042.

**Table 1.** Demographic, behavioral, and physiological data.

|  | Men | Women | p<sup>#</sup> | t<sup>#</sup> | Cohen's d |
| --- | --- | --- | --- | --- | --- |
| Number of subjects | 263 |  |  |  |  |
|  | 134 | 129 |  |  |  |
| Age [years] | 23.5±2.8 | 23.0±2.8 | p=0.123 | 1.55 | 0.19 |
| AQ | 2.20±0.44 | 2.18±0.43 | p=0.753 | 0.31 | 0.04 |
| Physical aggression | 1.78±0.57 | 1.40±0.46 | **p<0.001** | **5.95** | **0.74** |
| Verbal aggression | 2.66±0.56 | 2.74±0.60 | p=0.225 | -1.22 | -0.15 |
| Hostility | 2.23±0.73 | 2.25±0.62 | p=0.823 | -0.22 | -0.03 |
| Anger | 2.13±0.65 | 2.33±0.70 | **p=0.014** | **-2.47** | **-0.30** |
| Saliva testosterone concentration [pmol/l] | 212.3±63.5 (n=40) | 16.9±6.9 (n=36) | **p<0.001<sup>‡</sup>** | **19.61<sup>‡</sup>** | **4.39** |

Mean values, standard deviations, and p-values according to a two-sample t-test are given. The table summarizes the results of the German version of the Buss and Perry Aggression Questionnaire (AQ) (Buss and Perry, 1992; Herzberg, 2003), which comprises four subscales: anger, physical aggression, verbal aggression and hostility. Saliva testosterone concentration were determined by liquid chromatography tandem mass spectrometry (Keevil et al., 2014). <sup>#</sup>) two-sample t-test <sup>‡</sup>) two-sample t-test unequal variance

### Correlations of amygdala volume with aggression and testosterone level

We did not find any significant correlations between CM, SF or LB amygdala volume and the AQ score or any of its subscales. Effects of total intracranial volume, age, gender, and MRI scanner were removed by a regression prior to the correlation analysis. We observed no univariate whole-brain voxel-wise associations between gray matter and trait aggression at a corrected level. Of all amygdala nuclei groups, only the right CM amygdala showed a significant, negative correlation to the testosterone concentration in female subjects: □=−0.45 and p=0.006. We did not find any significant correlations between testosterone values and amygdala volume in male subjects (Table S1; supplementary materials). We did not find a correlation between whole-brain voxel-wise gray matter volume and testosterone concentrations in the male or female data sample.

### Structural covariance results

The centromedial (CMA), laterobasal (LBA), and superficial (SFA) amygdala nuclei groups showed distinct patterns of structural covariance (Figure 1; Table 2). The CMA had positive covariance with the basal ganglia, the LBA was associated with the rostral temporal gyrus, and the SFA showed positive covariance with a widespread network of brain regions comprising the insula, the anterior cingulate cortex (ACC) and the posterior cingulate cortex and adjacent precuneus.

**Figure 1.**
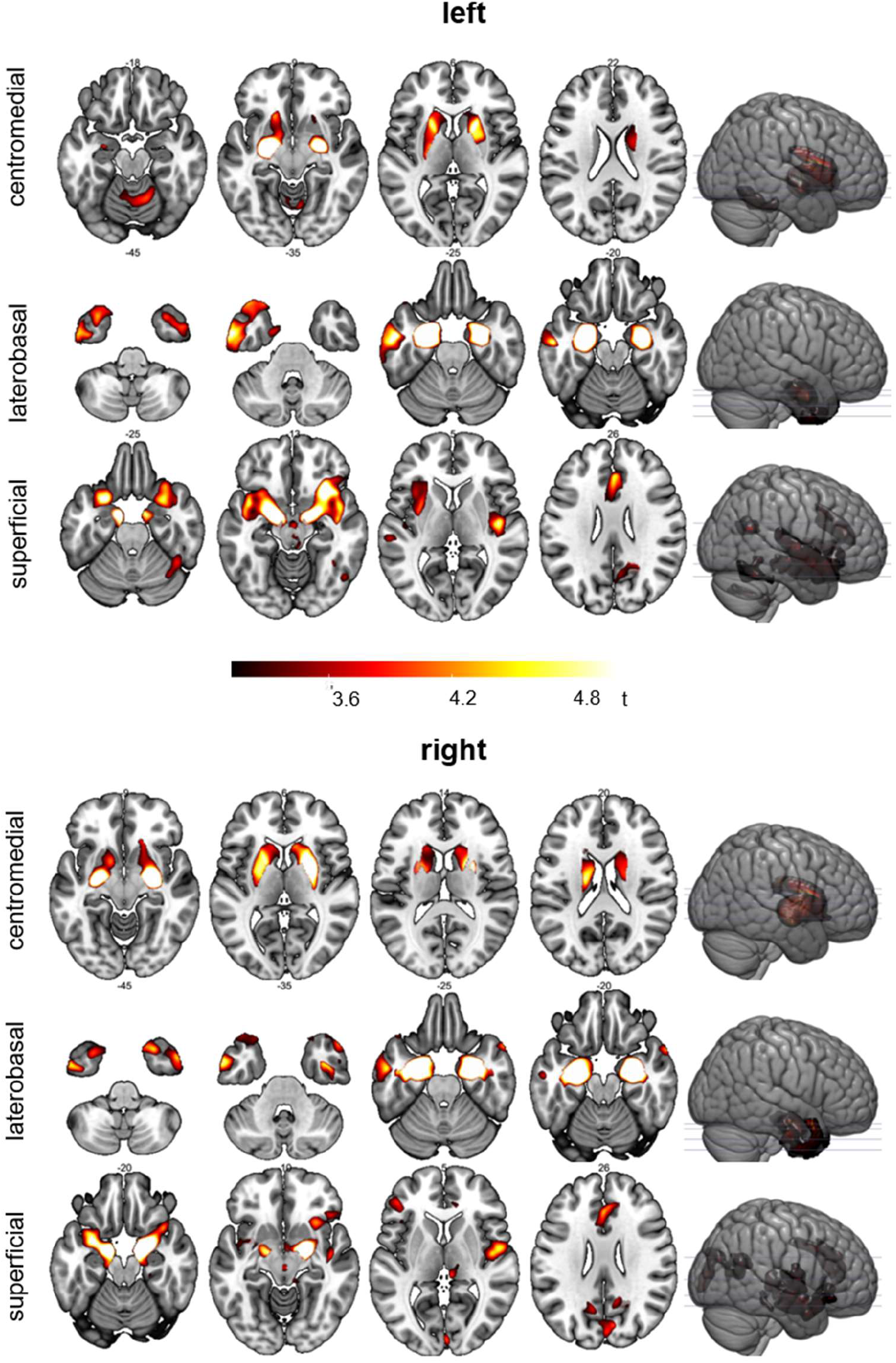
Structural covariance of amygdala subnuclei groups. Shown are the results for the centromedial (CMA, laterobasal (LBA) and superficial (SFA) amygdala nuclei groups (left and right hemispheres). A cluster-wise control of family-wise error was applied. The cluster defining threshold was p=0.001 and the 0.05 FWE corrected critical cluster size was k=713. The individual nuclei groups showed distinct patterns of structural covariance. The CMA volume was most strongly correlated to the basal ganglia volume, the LBA volume to the rostral temporal lobe and the SFA volume to a widespread network of brain regions comprising the insula, anterior cingulate cortex and the posterior cingulate cortex/precuneus.

**Table 2.** Structural covariance of the centromedial (a), laterobasal (b) and superficial (c) amygdala nuclei groups.

| *Brain region (at local maximum)* | *Hem* | *p (FWE) cluster* | *Cluster size* | *p (FWE) peak* | *T peak* | *MNI coord.[mm]* |
| --- | --- | --- | --- | --- | --- | --- |
| **a) centromedial amygdala** |  |  |  |  |  |  |
| **CMA left** |  |  |  |  |  |  |
| centromedial amygdala | L | 0.000 | 7900 | 0.000 | 30.38 | -22,-9,-8 |
| centromedial amygdala | R |  |  | 0.000 | 8.51 | -24,-8,-8 |
|  | R |  |  | 0.000 | 5.93 | 27,3,14 |
| *cerebellum* | R | 0.001 | 1793 | 0.331 | 4.14 | 8,-63,-12 |
|  |  |  |  | 0.335 | 4.14 | 12,-32,-26 |
|  |  |  |  | 0.417 | 4.06 | 15,-57,18 |
| **CMA right** |  |  |  |  |  |  |
| centromedial amygdala | R | 0.000 | 5677 | 0.000 | 25.56 | 27,-9,-9 |
|  |  |  |  | 0.000 | 6.79 | 27,3,14 |
|  |  |  |  | 0.000 | 6.78 | 20,2,9 |
| centromedial amygdala | L | 0.000 | 5306 | 0.000 | 7.77 | -24,-9,-8 |
|  |  |  |  | 0.005 | 5.28 | -20,0,10 |
|  |  |  |  | 0.016 | 4.99 | -18,9,4 |
| **b) laterobasal amygdala** |  |  |  |  |  |  |
| **LBA left** |  |  |  |  |  |  |
| laterobasal amygdala | L | 0.000 | 3098 | 0.000 | 38.16 | -26,-4,-24 |
| laterobasal amygdala | R | 0.001 | 1791 | 0.000 | 10.86 | 28,-3,-26 |
| rostral temporal lobe | L | 0.000 | 5264 | 0.002 | 5.48 | -56,-10,-22 |
| rostral temporal lobe | R | 0.023 | 875 | 0.021 | 4.92 | 48,3,-50 |
| **LBA right** |  |  |  |  |  |  |
| laterobasal amygdala | R | 0.000 | 4923 | 0.000 | 35.79 | 28,-3,-24 |
| rostral temporal lobe | R |  |  | 0.002 | 5.50 | 30,12,-51 |
| laterobasal amygdala | L | 0.000 | 3334 | 0.000 | 10.83 | -28,-6,-24 |
| rostral temporal lobe | L | 0.001 | 1737 | 0.006 | 5.2 | -62,0,-33 |
| rostral temporal lobe | L | 0.01 | 1058 | 0.176 | 4.35 | -28,10,-50 |
| **c) superficial amygdala** |  |  |  |  |  |  |
| **SFA left** |  |  |  |  |  |  |
| superficial amygdala | L | 16885 | 0.000 | 0.000 | 22.94 | -15,-4,-18 |
| superficial amygdala | R |  |  | 0.000 | 8.93 | 14,-6,-15 |
| insula | L |  |  | 0.001 | 5.55 | -44,8,-8 |
| insula | R |  |  | 0.002 | 5.45 | 46,-3,-4 |
| anterior cingulate cortex | R | 2098 | 0.000 | 0.039 | 4.77 | 2,33,22 |
| anterior cingulate cortex | L |  |  | 0.459 | 4.02 | -2,18,36 |
| cerebellum | L | 1333 | 0.000 | 0.273 | 4.21 | -24,-44,-48 |
| precuneus | R | 972 | 0.015 | 0.364 | 4.11 | 27,-52,22 |
| cerebellum | R | 1035 | 0.012 | 0.404 | 4.07 | 42,-52,-22 |
| **SFA right** |  |  |  |  |  |  |
| superficial amygdala | R | 5183 | 0.000 | 0.000 | 22.83 | 20,-3,-16 |
| superficial amygdala | L | 2623 | 0.000 | 0.000 | 7.72 | -16,-4,-16 |
| parahippocampus | R | 749 | 0.04 | 0.007 | 5.19 | 32,-30,3 |
| anterior cingulate cortex | L | 2178 | 0.000 | 0.032 | 4.82 | -15,28,32 |
| anterior cingulate cortex | R |  |  | 0.057 | 4.67 | 3,30,30 |
| inferior frontal gyrus | L | 1037 | 0.011 | 0.058 | 4.67 | -42,36,2 |
| insula | R | 1592 | 0.001 | 0.052 | 4.69 | 52,-4,6 |
| precuneus | R | 2908 | 0.000 | 0.292 | 4.19 | 16,-66,15 |

Statistical maps were assessed applying a cluster-wise FWE correction (cluster forming threshold $p=0.001$ ; 0.05 FWE-corrected critical cluster size $k=713$ ).

We tested for significant differences in structural covariance between amygdala nuclei groups. In figure 2 we show FWE-corrected (red; cluster-wise inference; cluster defining threshold p=0.001 and 0.05 FWE corrected critical cluster size k=713) and uncorrected data (blue; cluster-wise inference; uncorrected cluster probability p=0.05; k>314). Compared to the LBA and CMA, the SFA showed increased connectivity to the insula, the lateral and medial orbitofrontal cortex, inferior frontal gyrus, the anterior and posterior cingulate cortex, the precuneus and the angular gyrus at a corrected level (Figure 2; Table3). We did not detect increased connectivity for the LBA or CMA relative to the SFA nor significant differences in structural connectivity between LBA and CMA.

**Figure 2.**
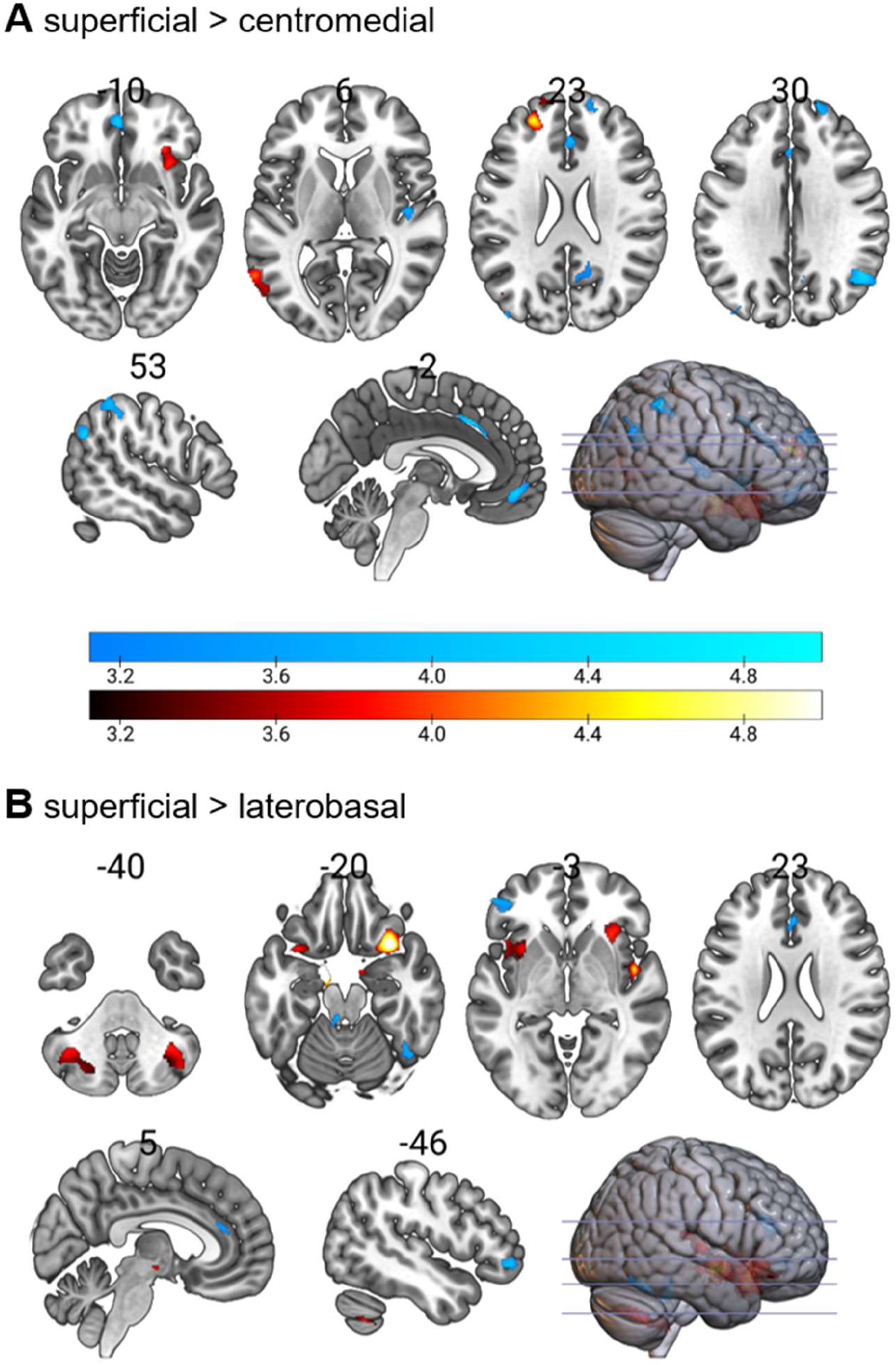
Differences in structural covariance between amygdala sublei groups. Show are differences between the superficial and the centromedial amygdala (A) and between the superficial and laterobasal amygdala (B). FWE-corrected (red; cluster-wise inference; cluster defining threshold p=0.001 and critical cluster size k=713) and uncorrected data (blue; cluster-wise inference; uncorrected cluster probability p=0.05; k>314) are shown. The SFA shows stronger structural covariance to the insula, the lateral and medial orbitofrontal cortex, the anterior and posterior cingulate cortex, the precuneus and the angular gyrus. We did not find significant differences between CM and LBA structural covariance.

**Table 3.**
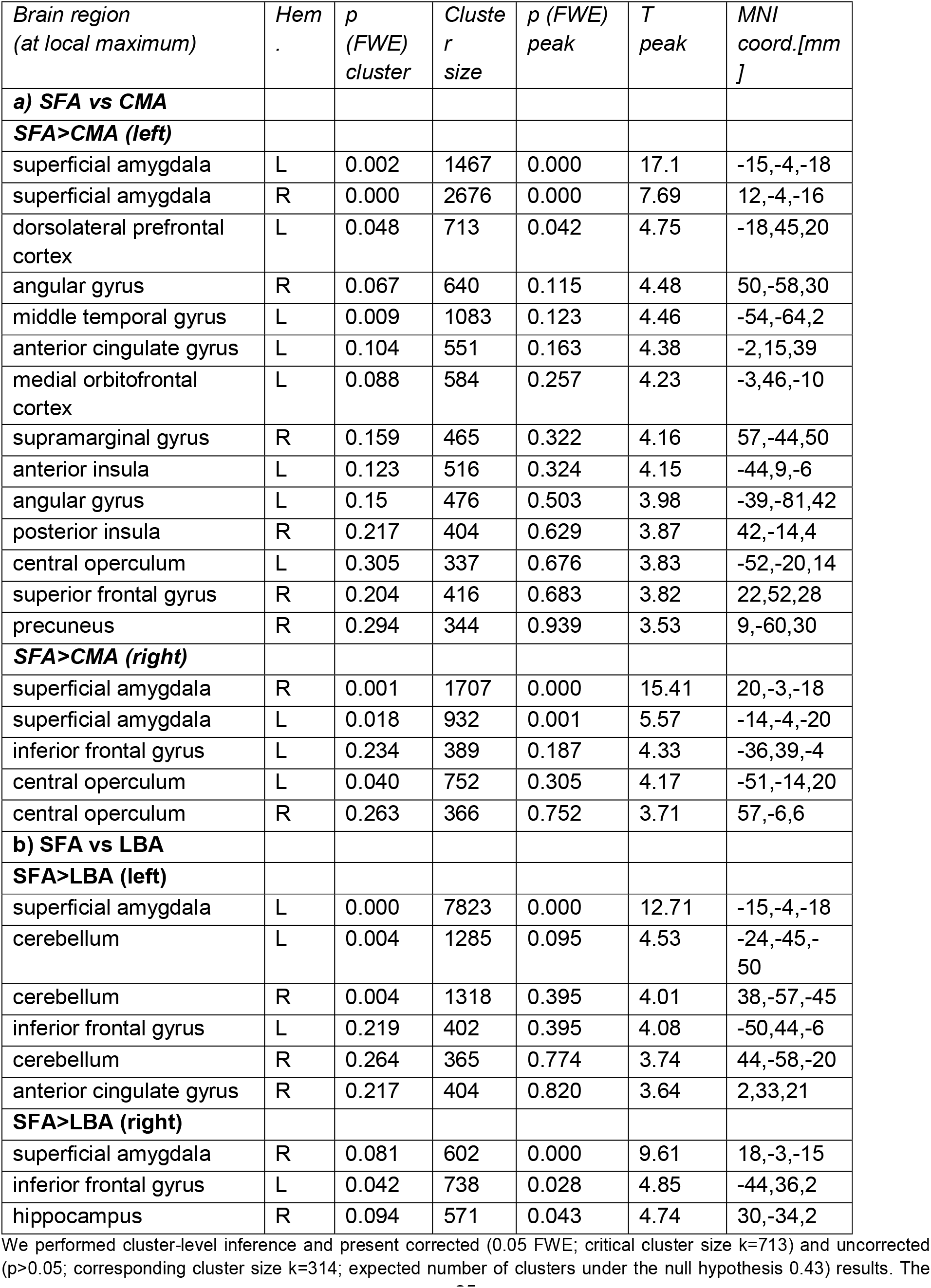
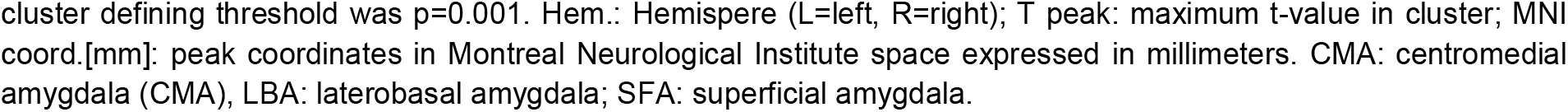
Differences in structural covariance between the amygdala subnuclei.

| *Brain region*<br>*(at local maximum)* | *Hem* | *p*<br>*(FWE)*<br>*cluster* | *Cluster*<br>*size* | *p (FWE)*<br>*peak* | *T*<br>*peak* | *MNI*<br>*coord.[mm*<br>*]* |
| --- | --- | --- | --- | --- | --- | --- |
| **a) SFA vs CMA** |  |  |  |  |  |  |
| **SFA>CMA (left)** |  |  |  |  |  |  |
| superficial amygdala | L | 0.002 | 1467 | 0.000 | 17.1 | -15,-4,-18 |
| superficial amygdala | R | 0.000 | 2676 | 0.000 | 7.69 | 12,-4,-16 |
| dorsolateral prefrontal cortex | L | 0.048 | 713 | 0.042 | 4.75 | -18,45,20 |
| angular gyrus | R | 0.067 | 640 | 0.115 | 4.48 | 50,-58,30 |
| middle temporal gyrus | L | 0.009 | 1083 | 0.123 | 4.46 | -54,-64,2 |
| anterior cingulate gyrus | L | 0.104 | 551 | 0.163 | 4.38 | -2,15,39 |
| medial orbitofrontal cortex | L | 0.088 | 584 | 0.257 | 4.23 | -3,46,-10 |
| supramarginal gyrus | R | 0.159 | 465 | 0.322 | 4.16 | 57,-44,50 |
| anterior insula | L | 0.123 | 516 | 0.324 | 4.15 | -44,9,-6 |
| angular gyrus | L | 0.15 | 476 | 0.503 | 3.98 | -39,-81,42 |
| posterior insula | R | 0.217 | 404 | 0.629 | 3.87 | 42,-14,4 |
| central operculum | L | 0.305 | 337 | 0.676 | 3.83 | -52,-20,14 |
| superior frontal gyrus | R | 0.204 | 416 | 0.683 | 3.82 | 22,52,28 |
| precuneus | R | 0.294 | 344 | 0.939 | 3.53 | 9,-60,30 |
| **SFA>CMA (right)** |  |  |  |  |  |  |
| superficial amygdala | R | 0.001 | 1707 | 0.000 | 15.41 | 20,-3,-18 |
| superficial amygdala | L | 0.018 | 932 | 0.001 | 5.57 | -14,-4,-20 |
| inferior frontal gyrus | L | 0.234 | 389 | 0.187 | 4.33 | -36,39,-4 |
| central operculum | L | 0.040 | 752 | 0.305 | 4.17 | -51,-14,20 |
| central operculum | R | 0.263 | 366 | 0.752 | 3.71 | 57,-6,6 |
| **b) SFA vs LBA** |  |  |  |  |  |  |
| **SFA>LBA (left)** |  |  |  |  |  |  |
| superficial amygdala | L | 0.000 | 7823 | 0.000 | 12.71 | -15,-4,-18 |
| cerebellum | L | 0.004 | 1285 | 0.095 | 4.53 | -24,-45,-50 |
| cerebellum | R | 0.004 | 1318 | 0.395 | 4.01 | 38,-57,-45 |
| inferior frontal gyrus | L | 0.219 | 402 | 0.395 | 4.08 | -50,44,-6 |
| cerebellum | R | 0.264 | 365 | 0.774 | 3.74 | 44,-58,-20 |
| anterior cingulate gyrus | R | 0.217 | 404 | 0.820 | 3.64 | 2,33,21 |
| **SFA>LBA (right)** |  |  |  |  |  |  |
| superficial amygdala | R | 0.081 | 602 | 0.000 | 9.61 | 18,-3,-15 |
| inferior frontal gyrus | L | 0.042 | 738 | 0.028 | 4.85 | -44,36,2 |
| hippocampus | R | 0.094 | 571 | 0.043 | 4.74 | 30,-34,2 |

We performed cluster-level inference and present corrected (0.05 FWE; critical cluster size $k=713$ ) and uncorrected ( $p>0.05$ ; corresponding cluster size $k=314$ ; expected number of clusters under the null hypothesis 0.43) results. The
cluster defining threshold was $p=0.001$ . Hem.: Hemisphere (L=left, R=right); T peak: maximum t-value in cluster; MNI coord.[mm]: peak coordinates in Montreal Neurological Institute space expressed in millimeters. CMA: centromedial amygdala (CMA), LBA: laterobasal amygdala; SFA: superficial amygdala.

### Relation of structural covariance with aggression and testosterone

We tested whether structural covariance of each amygdala nucleus group was modulated by trait aggression, i.e., the interaction between each nucleus’ volume and AQ score. The left SFA volume showed an increasing correlation with the left dorsal anterior insula (dAI; peak t=4.80, x=−26 mm, y=26 mm, z=12 mm, k=734) for higher aggressiveness (Figure 3a). In Figure 3b we visualized this effect and plotted the correlation between the SFA and the dAI volume subdividing the whole data sample into three groups of increasing trait aggression. For subjects with low AQ scores (mean AQ 1.52; range 1 to 1.86) we observed a significant negative correlation (rho = −.25). For the medium aggression sample (mean AQ 2.15; range 1.86 to 2.5) the correlation turned to a significantly positive value (rho = .21) and continued to rise in the high aggression sample (mean AQ 3.00; range 2.5 to 4.17; rho = .44). The correlation held across all subscales (physical aggression, verbal aggression, hostility, and anger) of the aggression questionnaire (Figure 3c).

**Figure 3.**
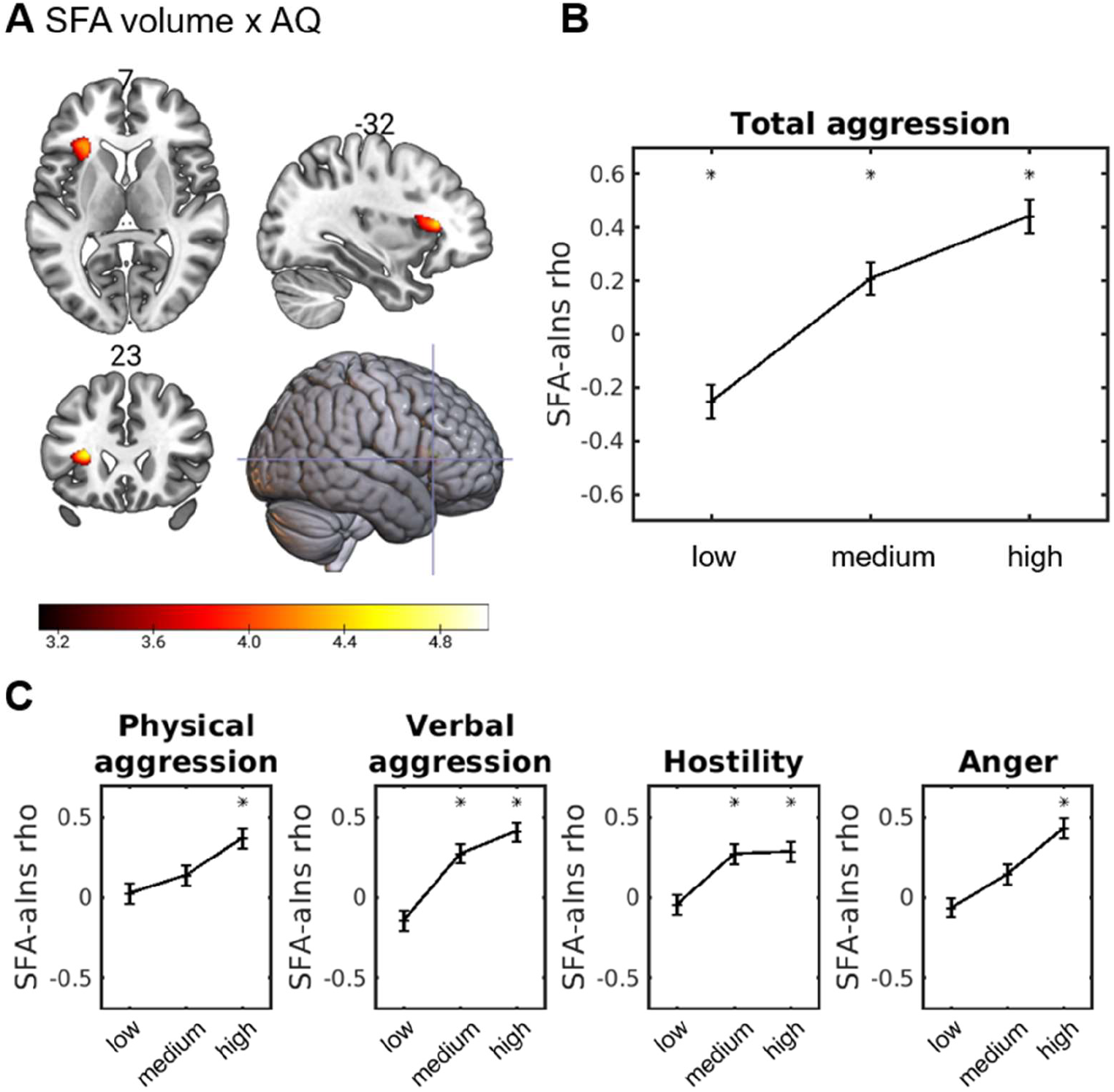
Structural covariance of amygdala and trait aggression. The left dorsal anterior insula (dAI) showed a significant interaction between the left superficial amygdala (SFA) volume and trait aggression (A). A cluster-wise FWE correction was applied (p<0.05 corrected). This effect is visualized in B. The data sample was subdivided into three equally large groups of subjects with increasing trait aggression. In each group the correlation between the dAI and SFA volume was calculated. The same approach was used for all four subscales of the AQ questionnaire separately (C).

The right LBA showed a decreasing correlation with the right dorsolateral prefrontal cortex (dlPFC; peak t=4.43, x=34 mm, y=39 mm, z=46 mm, k=789) in subjects with higher trait aggressiveness (Figure 4a). For the low and medium aggressive subsample, the correlation was positive (albeit not significant) but significantly negative for the high aggression group (Figure 4b). Figure 4c indicates a tendency for this effect to be mainly driven by verbal aggression and anger subscales. There were no aggression-dependent associations between the CMA and the rest of the brain at a corrected level.

**Figure 4.**
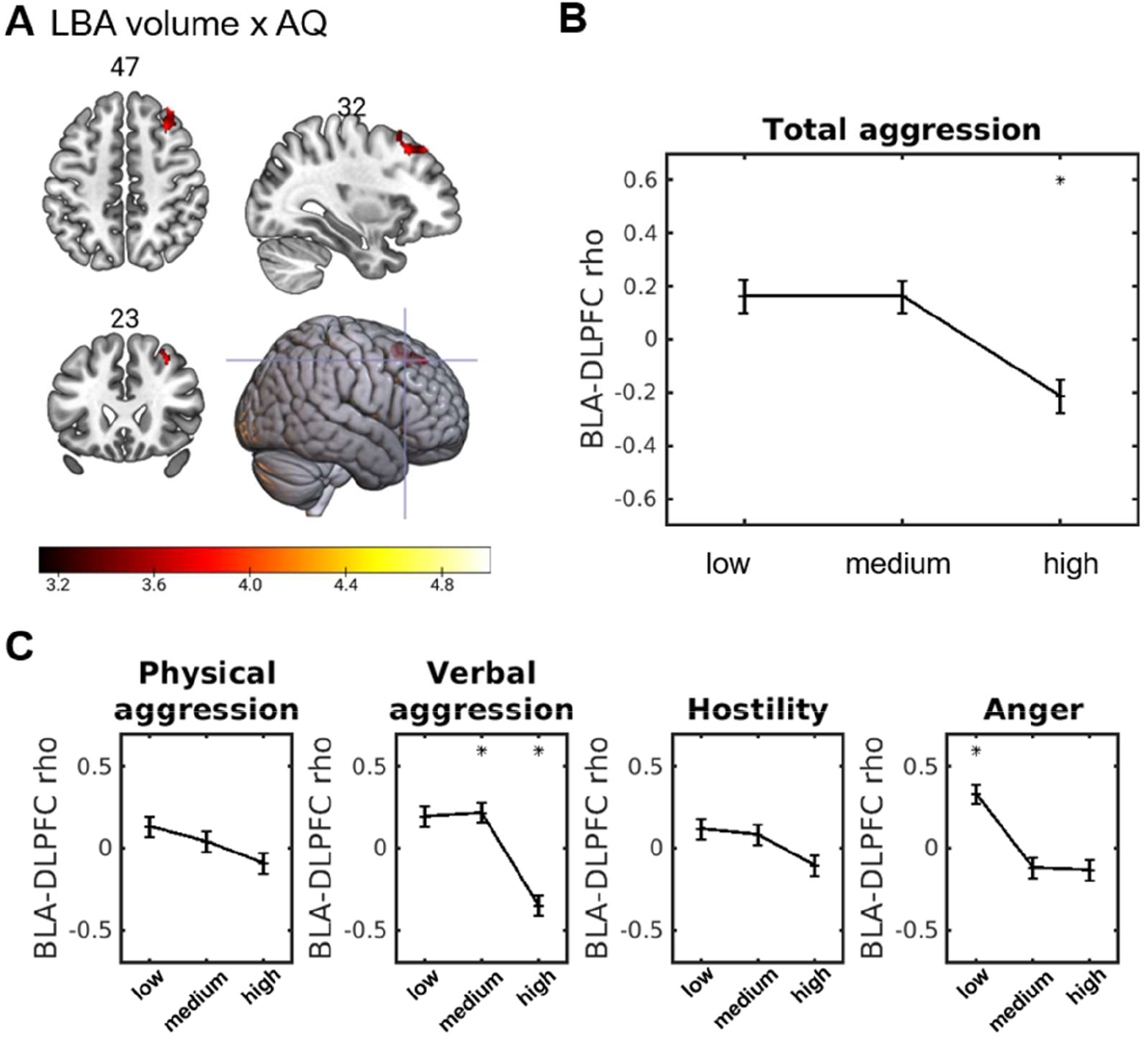
Structural covariance of amygdala and trait aggression. Aggression-dependent correlation between the right dorsolateral prefrontal cortex (DLPFC) and the laterobasal amygdala (LBA) volume (A). B In the high aggression group, the correlation was significantly negative. For the low and medium aggression group the correlation was positive but not significant (p>0.05). For the different subscores we obtained consistent results suggesting a positive correlation for low and medium aggression and an anti-correlation for high aggression (C).

With respect to testosterone, we found an increasing correlation between right CM amygdala and right medial frontal gyrus (MFG) volume for increasing testosterone levels (Figure 5a; cluster peak t=4.84, x=14 mm, y=39 mm, z=36 mm, cluster size k=983; cluster-level FWE-corrected p=0.05) in male subjects. We subdivided the male data sample (n=40) into three groups of increasing testosterone levels and determined the correlation between CMA and MFG volume in each group (Figure 5b). For subjects with high testosterone (mean testosterone level 284.0±38.1 pmol/l; range 241.0 to 366.3 pmol/l) we observed a significant positive correlation (Pearson’s □=0.77; p=0.0013). For low (mean 147.3±34.1 pmol/l; range 96.9 to 192.2 pmol/l; □=−0.41; p=0.1642) and medium mean (215.5±12.25 pmol/l; range 192.9 to 236.5 pmol/l; □=−0.19; p=0.5132) levels of testosterone the correlation was negative but not significant. Women did not show a similar effect (Figure 5c).

**Figure 5.**
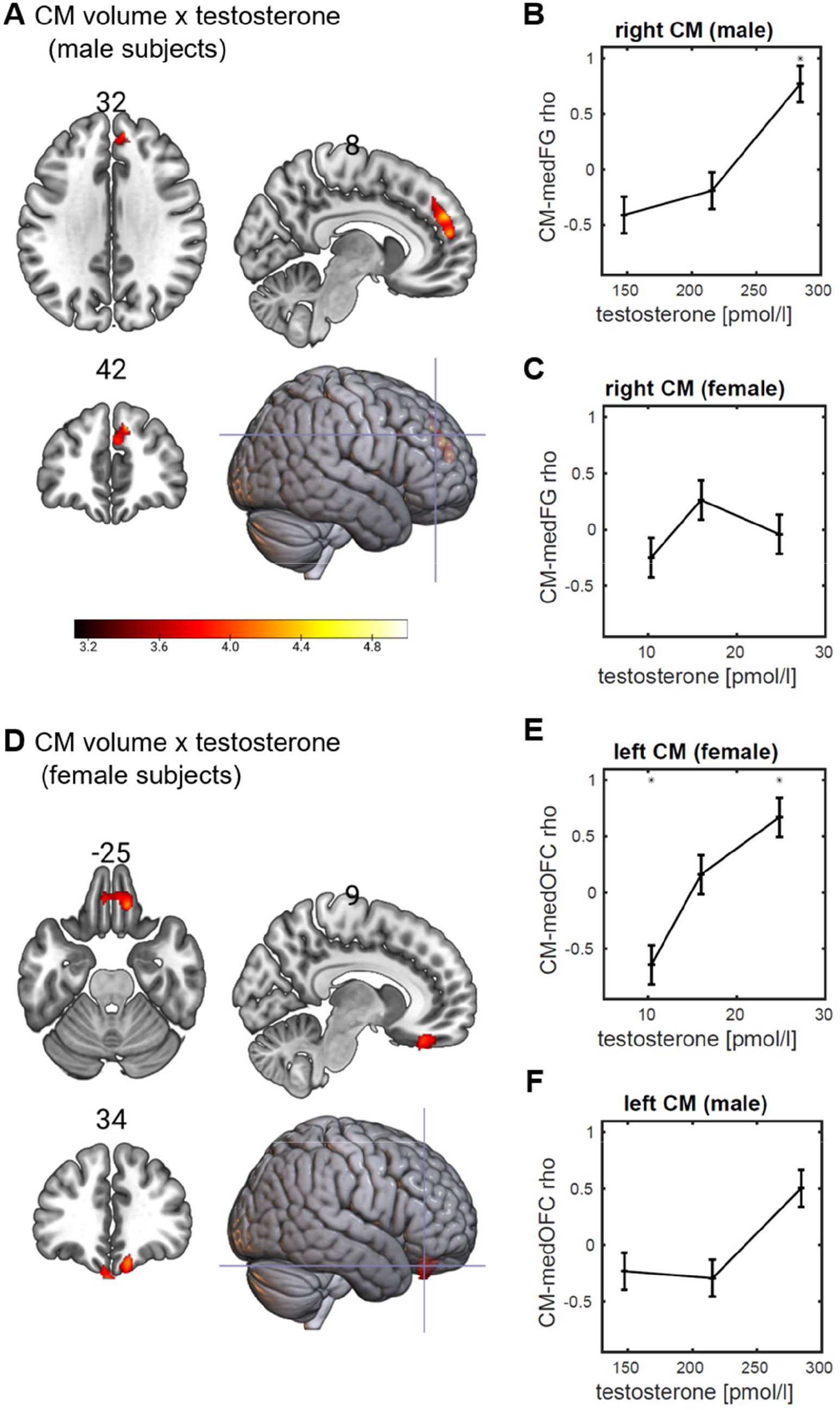
Amygdala structural covariance and testosterone level. Interaction between saliva testosterone concentrations and centromedial amygdala (CMA) gray matter volume. (A) Male subjects showed a significant positive interaction in the medial frontal gyrus (medFG). For high testosterone levels the correlation between CMA and medFG volume was positive and turned negative for low testosterone (B). This effect was not observed in female subjects (C). (D) Female subjects showed a significant positive interaction in the medial orbitofrontal gyrus (medOFC). Low testosterone was related to a negative correlation between CMA and medOFC volume and high testosterone level to a positive correlation (E). There was a tendency for male subjects to show a similar pattern. Statistic images were assessed for cluster-wise significance and 0.05 FWE-corrected.

In female subjects (n=36), increasing testosterone levels were linked to a stronger correlation between CM amygdala and medial orbitofrontal cortex (mOFC) gray matter volume (Figure 5d; cluster peak t=4.74; MNI coordinates x=−4mm, y=33mm, z=−34mm; cluster size k=910; cluster-level FWE-corrected p=0.05). For low testosterone (mean testosterone level 10.4±2.7 pmol/l; range 5.0 to 13.1 pmol/l) women (n=12 per group) showed a significant negative correlation between these two brain regions (□=−0.64; p=0.0238; Figure 5e). High testosterone (24.8±5.1 pmol/l; range 19.2 to 34.2 pmol/l) was linked to a significantly positive correlation (□=−0.67; p=0.0175). For medium testosterone levels (16.0±1.7 pmol/l; range 13.5 to 18.4 pmol/l) the correlation was not significant (□=0.16; p=0.6210). Male subjects did not show a significant interaction between testosterone level and amygdala volume in the mOFC. However, the ROI-based analysis showed a positive but not significant correlation in the mOFC for high testosterone in male subjects (Figure 5f; □=0.50; p=0.0674) which might indicate a similar behavior in men compared to women and which should be tested in a future analysis using an independent data set. Summarizing, although the data speak for an impact of testosterone level on amygdala dependent structural covariance, the implicated brain regions differ from the networks related to trait aggression as reported above.

Our results could be reproduced in each of the three data sets from different MRI scanners (Philips Achieva, Philips Ingenia and Siemens Skyra). We show this as an example in Figure S1 (supplementary materials). Controlling for total intracranial volume, age, and gender effects, we found a higher correlation between SFA and anterior insula volume for increasing trait aggression in all scanner models.

## Discussion

We investigated whether trait aggression in humans is linked with structural covariance of the three main amygdala nuclei groups in a large sample of healthy young students. Relatively more aggressive subjects evinced greater structural covariance between superficial amygdala (SFA) and the dorsal anterior insula (dAI). Aggressive individuals also displayed a reduced volumetric correlation between laterobasal amygdala (LBA) and dorsolateral prefrontal cortex. We hence delineated neurostructural patterns that can potentially serve as a marker of aggressiveness in non-clinical populations. Our analysis of testosterone modulated amygdala covariance revealed a positive interaction between testosterone level and CM amygdala volume in the medial PFC in men. Women showed a positive interaction between testosterone level and CM amygdala volume in den medial OFC.

### Trait aggression and endogenous testosterone

We used the German version of the Buss and Perry Aggression Questionnaire (AQ) (Buss and Perry, 1992; Herzberg, 2003) to assess trait aggression in our participants. Compared to Herzberg (2003), our participants tended to report lower levels of aggression. This effect was even stronger for male participants, which mitigated the expected gender differences in aggression in our sample. The only subscale where men showed significantly higher scores than women was physical aggression but their score was still lower than what was reported by Herzberg (2003). Self-reported anger was significantly higher in women compared to men while it was in good agreement with Herzberg (2003). The most likely explanation for decreased aggression scores compared to Herzberg (2003) were differences in the composition of the samples. Our sample consisted of college students (convenience sample) while Herzberg (2003) recruited a more representative sample and hence obtained a wider value distribution.

Physical aggression was positively correlated to saliva testosterone concentrations in men. Combining the two data sets of male and female subjects, we found a significant correlation between aggression (AQ) and testosterone (z-transformed within gender prior to the analysis). Our results are in good agreement with a recent meta-analysis by Geniole et al. (2020) who reported a positive correlation between baseline testosterone and human aggression. Also, in line with the meta-analysis, this effect was significant in men, but not women, when the correlations were computed separately for each group.

### Structural covariance of amygdala subregions

We first inspected the structural network of the amygdala subregions, which showed distinct patterns of structural covariance. Centromedial amgydala (CMA) volume was associated with the basal ganglia and motor regions in the cerebellum, perhaps reflecting a recently described intrinsic network thought to underlie motor preparation (Toschi et al., 2017). Interestingly, this pattern of structural covariance is similar to functional connectivity derived from resting-state fMRI data (Roy et al., 2009) and to coactivation patterns determined by meta-analyses (Bzdok et al., 2013). The LBA was strongly connected to the rostral temporal lobe which was also reported by Roy et al. (2009), who suggested that this connection may facilitate associative learning processes (Schoenbaum et al., 2000). The SFA showed the most widespread network of structural covariance, encompassing the precuneus, dorsal cerebellum, dorsolateral prefrontal cortex, anterior insula (AI) and anterior cingulate cortex (ACC). Contrasting SFA covariance to the covariance patterns of the CMA and LBA, we identified regions of the salience network (ACC and AI) and of the mentalizing network (angular and supramarginal gyrus, anterior cingulate cortex, and medial frontal cortex, precuneus). This fits to the view that the SFA is involved in processing social information (Bzdok et al., 2013; Bzdok et al., 2012; Goossens et al., 2009; Haruno and Frith, 2010). Moreover, Haruno and Frith (2010) reported that prosocial value orientation was related to the superficial nuclei group. We conclude that the observed structural covariance patterns partially reproduce well-known functional networks. Notably, there was no negative correlation between amygdala volume and the rest of the brain.

### Trait aggression and structural covariance of amygdala subregions

We subsequently tested whether structural covariance patterns of amygdala nuclei were modulated by trait aggression. There was enhanced volume covariance between superficial amygdala (SFA) and dorsal anterior insula (dAI) as a function of aggressiveness. Along with the ACC, both SFA and dAI are part of the salience network, which has been previously documented at both the structural and functional level (Seeley et al., 2009; Seeley et al., 2007) and which we also observed in the present dataset. The anterior insula and the inferior frontal gyrus have also been shown to be involved in processing socially salient information (Carr et al., 2003; Göttlich et al., 2017). Although structural covariance between ACC and insula increases gradually during development, it is only in late adolescence (i.e. 16-18 years of age) that these regions first show a robust positive association with amygdala volume (Zielinski et al., 2010). Across studies, both amygdala and anterior insula display lower volume in chronically aggressive youth (Rogers and De Brito, 2016). Our results expand on this and show that increased structural covariance between SFA and dAI is linked with an enhanced propensity for antagonistic behavior. This might be due to the involvement of both regions in experiencing frustration and negative affect in social situations (Buades-Rotger et al., 2016b; Dambacher et al., 2014; Yu et al., 2014). Notably however, trait aggression did not correlate with grey matter volume per se in any of these regions in the current study.

In addition, we observed a negative correlation between LBA and dorsolateral prefrontal cortex (dlPFC) volume covariance and trait aggression. Lesions in dlPFC are associated with more positive implicit attitudes toward violence (Cristofori et al., 2016) and have been generally linked with deficits in executive function (Szczepanski and Knight, 2014), an established predictor of aggressive behavior in at-risk populations (Micai et al., 2015; Tonnaer et al., 2016). Paralleling our results, a recent investigation reported reduced functional LBA-dlPFC coupling at rest in veterans with anger and aggression problems relative to non-aggressive combat-exposed controls (Varkevisser et al., 2017). This speaks for an overlap between functional and structural amygdala networks involved in the regulation of aggressive behavior, and potentially indicates that low LBA-dlPFC structural covariance signals a lack of modulatory crosstalk between these two regions. Future studies should elucidate the precise anatomical pathways underlying this aggression-related link, as there are no strong, direct reciprocal fibers connecting BLA and dlPFC (Abivardi and Bach, 2017).

It should also be noted that we observed no univariate whole-brain associations between gray matter and trait aggression at a corrected level. We also did not observe any significant correlations between CM, SF and BL amygdala volume and trait aggression or any of the AQ subscales. In the latter analysis we extracted the mean gray matter volume in the different amygdala nuclei in contrast to a voxel-wise approach as in the whole-brain analysis. Although caution must be exercised regarding negative findings, a study with a comparable but smaller sample than the present one did yield a negative correlation between ventral prefrontal volume and physical aggressiveness (Chester et al., 2017). Pending independent replications of these and the current findings, the present data suggest that normal variation in aggressiveness might be better explained by interregional structural correlations than by the volume of single brain areas. It should be stressed that also methodological differences must be considered to explain the inconsistent findings. For instance, Chester et al. (2017) applied the Brief Aggression Questionnaire (Webster et al., 2014) to assess trait aggression and did not correct for total intracranial volume (TIV) and age in contrast to the current study. As there is no widely accepted gold standard for the analysis of VBM data, this hinders and limits the comparability across studies.

### Testosterone-dependent structural covariance of amygdala subregions

Women showed a testosterone dependent correlation between centromedial amygdala (CMA) and medial orbitofrontal cortex volume. Low testosterone was associated with a negative correlation between these brain regions and higher levels of testosterone were associated with a positive correlation. There was a trend in male subjects to show a similar pattern. The orbitofrontal cortex and the amygdala are reciprocally connected, and both are crucial brain regions for processing emotions (Abivardi and Bach, 2017) with the orbitofrontal cortex involved in emotional regulation and impulse control (Beyer et al., 2015; Dixon et al., 2017). An increased correlation between CMA and orbitofrontal volume might be related to increased connectivity between these two regions (Alexander-Bloch et al., 2013) mediating top-down regulation of the amygdala promoted by high baseline testosterone in women. This assumption is supported by recent evidence for decreased amygdala reactivity to angry faces with increasing endogenous testosterone levels in the same female participants (Buades-Rotger et al., 2016b). Interestingly, amygdala and medial orbitofrontal cortex effective connectivity was reduced when subjects were confronted with angry faces and this in turn led to higher aggression. Taken together, the two observations led to the conclusion that the OFC is downregulating the amygdala when being confronted with threat or being provoked and that this effect is stronger for high baseline testosterone. One of the structural underpinnings of this mechanism might be a stronger structural connectivity between medial OFC and amygdala for high baseline endogenous testosterone. This effect is consistent with the fear-reducing and strategically prosocial influence of testosterone on human social behavior, particularly in women (Eisenegger et al., 2011).

On the other hand, men showed an increasing correlation between centromedial amygdala (CMA) and medial prefrontal volume for increasing endogenous testosterone. This observation might be explained by a regulatory influence on the amygdala, which is promoted by testosterone. Both ventral and dorsal aspects of the medial prefrontal cortex show increased connectivity with the amygdala during effortful emotion regulation (Scharnowski et al., 2020). Furthermore, greater anatomical coupling of both regions with the amygdala predicts lower reactivity to emotional stimuli in the latter structure (Goetschius et al., 2019). Nguyen et al. (2016) investigated testosterone-dependent structural covariance in children, adolescents, and young adults. The sample comprised male and female subjects and the authors found no sex-related effects on testosterone-related structural covariance. The mean age was significantly lower compared to our study (13.5 years; SD 3.6 years). They reported a testosterone-dependent correlation between the amygdala volume and cortical thickness of the medial PFC, with a positive correlation for lower levels of testosterone but negative for higher concentrations. In contrast to our study, Nguyen et al. (2016) investigated covariance between amygdala volume and cortical thickness, which limits comparability between the studies (Carmon et al., 2020). Furthermore, in the present study the amygdala nuclei showed different covariance patterns, questioning the approach of using the whole amygdala as seed region. The main cause for the observed differences is likely to lie in the age disparity of the test subjects. Indeed, there is evidence for changing structural covariance networks across the human lifespan with largely non-linear alterations between the ages 5 and 18 years (Alexander-Bloch et al., 2013). Note that the present male and female subsamples were acquired at different sites, and we thus refrained from combing the two samples as we cannot disentangle the effects from MRI scanner and gender. The causes of possible gender differences are an interesting field for future targeted studies.

Notably, regions that showed structural covariance with the amygdala which was modulated by either testosterone or aggression did not overlap. This observation might reflect the weak link between testosterone and aggression and supports the notion of a more complex role of testosterone in human behavior rather than the simple assumption that testosterone only increases aggressiveness. Examples are the implication of testosterone in social status and in prosocial behavior (Buades-Rotger et al., 2016b; Eisenegger et al., 2011).

### Limitations

The cross-sectional nature of our data precludes testing for the developmental substrate of the observed effects. Furthermore, we could not establish clear links between brain structure and function because subjects performed different tasks in the scanner. However, our relatively large and homogeneous sample supports the robustness of our findings, at least within the confines of the population under study. Datasets such as the Adolescent Brain Cognitive Development (ABCD) study hold promise to help answer some of the questions left open by our results (Casey et al., 2018). One should also note that the number of subjects for whom testosterone levels were measured was limited in our current sample which limits the conclusions we can derive from it.

### Conclusions

We found that within healthy young persons, aggressive disposition was positively related to structural covariance between the superficial amygdala and the dorsal anterior insula, two regions that become increasingly correlated during development (Zielinski et al., 2010) and which are relatively smaller in pathologically aggressive youth (Rogers and De Brito, 2016). Moreover, aggressive subjects showed a weaker correlation between the volume of the laterobasal amygdala and dorsolateral prefrontal cortex, mimicking the reduced functional coupling between these two areas that has also been reported in veterans with elevated anger and aggression (Varkevisser et al., 2017). Further, the centromedial amygdala showed enhanced testosterone-dependent covariance with the medial prefrontal cortex in men and with the orbitofrontal cortex in women. In contrast to previous accounts, trait aggression and testosterone related amygdala dependent structural covariance thus did not converge in joint brain networks. This speaks against a major role of testosterone in driving the observed structural covariance patterns in high trait aggression. Our study uncovers a neurostructural phenotype that might prove valuable in understanding interindividual differences in aggressive behavior.

## Acknowledgements

This study was funded by the German Science Foundation (grant number KR3691/5-1). MBR is supported by a German Science Foundation fellowship (BU3756/1-1). We are grateful to Susanne Schellbach and Christian Erdmann for their help with the data acquisition. The authors report no conflicts of interest.

