## Supplementary Information for "Structural covariance of amygdala subregions is associated with trait aggression and endogenous testosterone in healthy individuals"

**Supplementary materials**

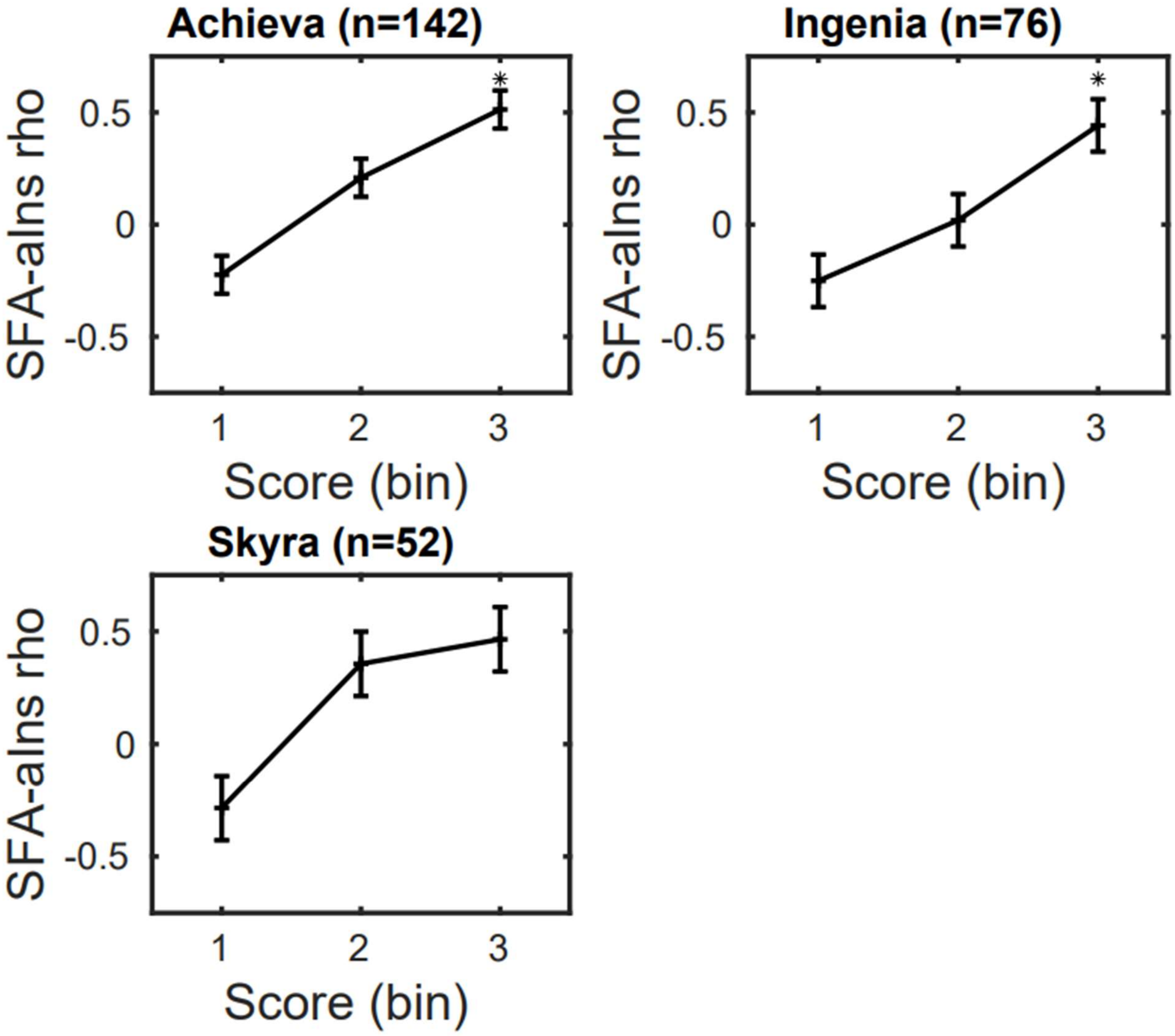


Figure S1: Correlation between the superficial amygdala subgroup and anterior insula volume as a function of increasing trait aggression for the three different datasets separately. The results of all three different subsets are compatible.

**Table S1. Correlation (Pearson's rho) between CM, LB, and SF amygdala volume and testosterone.**

| | Male subjects (n=40) | | Female subjects (n=36) | |
|---|---|---|---|---|
| | rho | p | rho | p |
| CM left | -0.11 | 0.511 | -0.02 | 0.906 |
| CM right | 0.01 | 0.967 | **-0.45** | **0.006** |
| LB left | -0.02 | 0.892 | 0.00 | 0.989 |
| LB right | -0.06 | 0.723 | -0.19 | 0.265 |
| SF left | 0.07 | 0.669 | -0.04 | 0.811 |
| SF right | -0.02 | 0.914 | -0.19 | 0.263 |
